# Incorporating Hierarchical Information into Multiple Instance Learning for Patient Phenotype Prediction with scRNA-seq Data

**DOI:** 10.1101/2025.02.10.637389

**Authors:** Chau Do, Harri Lähdesmäki

**Affiliations:** Aalto University

## Abstract

Multiple Instance Learning (MIL) provides a structured approach to patient phenotype prediction with single-cell RNA-sequencing (scRNA-seq) data. However, existing MIL methods tend to overlook the hierarchical structure inherent in scRNA-seq data, especially the biological groupings of cells, or cell types. This limitation may lead to suboptimal performance and poor interpretability at higher levels of cellular division. To address this gap, we present a novel approach to incorporate hierarchical information into the attention-based MIL framework. Specifically, our model applies the attention-based aggregation mechanism over both cells and cell types, thus enforcing a hierarchical structure on the flow of information throughout the model. Across extensive experiments, our proposed approach consistently outperforms existing models and demonstrates robustness in data-constrained scenarios. Moreover, ablation test results show that simply applying the attention mechanism on cell types instead of cells leads to improved performance, underscoring the benefits of incorporating the hierarchical groupings. By identifying the critical cell types that are most relevant for prediction, we show that our model is capable of capturing biologically meaningful associations, thus facilitating biological discoveries.

## 1 Introduction

Single-cell RNA sequencing (scRNA-seq) has the potential to transform personalized medicine by providing high-resolution, disease-specific signatures (He et al., 2021). This source of data can improve the prediction of patient-level characteristics, including disease phenotypes and clinical outcomes. During prediction, relevant cells and cell types can be identified, hence supporting biological discoveries (Xiong, Bekiranov, and Zhang, 2023).

In recent years, several methods have been developed for the prediction of patient-level characteristics using scRNA-seq data. Notably, multiple instance learning (MIL) emerges as a promising framework due to its ability to structurally model cellular heterogeneity (Engelmann et al., 2024). Specifically, the attention-based aggregation technique has reached state-of-the-art performance as it can model the influence of individual cells on the sample-level prediction in the form of attention weights (Ilse, Tomczak, and Welling, 2018). The integration of attention-based MIL with a generalized linear mixed model (GLMM) also reduces the problem of the low signal-to-noise ratio in scRNA-seq data and improves robustness (Engelmann et al., 2024).

However, existing MIL models do not account for the hierarchical structure of scRNA-seq data, especially the biological groupings of cells (i.e. cell types). This possibly leads to suboptimal performance, as these models might fail to capture the similarities of cells within a cell type in predicting the phenotype. In addition, the attention-based aggregation technique only offers interpretability at the cell level and not at the cell type level, which makes it difficult to draw associations between cell types and phenotypes. A common workaround is to approximate the cell type contribution as an average of the cell contributions. However, this approach does not account for the heterogeneity of cells within a cell type and therefore might not work well when the proportion of influential cells within a cell type is small.

In this work, we introduce an approach to incorporate hierarchical information into the attention-based MIL framework. Our approach employs existing cell type annotations to perform a step-by-step aggregation procedure, first over cells and subsequently over cell types, enforcing a hierarchical structure on the flow of information throughout the model. Specifically, we propose two aggregation strategies, represented in the Cell Type Attention (CTA) model and the Hierarchical Attention (HA) model. The CTA model combines mean-pooling of cells and attention-based aggregation of cell types, thereby reducing the complexity of scRNA-seq data yet still accounting for the heterogeneity of cell types in predicting the label. Meanwhile, the HA model performs attention-based aggregation of both cells and cell types, thus additionally capturing the heterogeneity of cells in a cell type. The sample-level predictions of both models can be decomposed into a weighted sum of either cell-level or cell type-level contributions, hence providing interpretability at both levels.

We benchmark our proposed models against four state-of-the-art models for patient phenotype prediction with scRNA-seq data. Across different disease phenotype prediction and clinical outcome modeling tasks, our models consistently achieve a competitive performance, underscoring the benefits of implement-ing a hierarchical design in a MIL framework.

## 2 Related Work

### 2.1 Multiple Instance Learning (MIL)

MIL refers to a specific setting of weakly supervised learning where data is organized into collections termed *bags*, each containing several *instances*. Note that the number of instances in each bag can vary. The key premise of MIL is that only the bag-level labels are known, and the instance-level labels are unknown. The task of MIL models is, therefore, to predict labels of bags using features of their instances. Given this setup, the problem of predicting phenotypes using scRNA-seq data naturally falls into the scope of MIL, in which bags are the samples from patients/donors, and the instances are the cells.

Ilse, Tomczak, and Welling, 2018 introduced the attention-based aggregation technique, which models the bag-level representation as an attention-weighted sum of the instance-level representations. The attention weights are modeled as a transformation of the instance-level representations, followed by a softmax activation over the instances. As the attention weights can be interpreted as the relative importance of instances, this aggregation technique allows MIL models to capture the cellular heterogeneity of scRNA-seq data and also provides interpretability at the single cell level.

Building upon this work, Engelmann et al., 2024 proposed MixMIL, a MIL model that integrates the attention-based aggregation technique with a generalized linear mixed model (GLMM). Specifically, MixMIL aggregates the cell representations into the sample representation with the attention technique, then models the dependency between the sample representation and the label with a GLMM. Notably, the attention function is a *shallow* function, consisting of a linear transformation of the cell representation and a softmax over the cells. This design allows MixMIL to simultaneously capture cellular heterogeneity with attention-based MIL and maintain robustness with a GLMM, combining the strengths of both frameworks.

Aside from modeling the relative importance of cells, the attention technique can also be used to capture the interactions between cells. Mao, Lin, Wong, et al., 2024 introduced ScRAT, a transformer-based model for patient phenotype prediction using scRNAseq data. Briefly, the attention mechanism in transformers iteratively updates the representation of a cell as a weighted sum of the representations of all other cells and of itself, hence capturing the influence of cells on one another.

The structured nature of MIL also allows for the incorporation of domain knowledge into the derivation of attention weights. Building upon the assumption that a prior distribution of the instance-level label is available for each instance, Hajj, Hubin, Kanduri, et al., 2024 proposed several strategies for prior knowledge incorporation, with the two most successful being attention modulation (AM) and attention training (AT). Here, the attention weight is formulated with sigmoid activation instead of softmax to represent the probability of an instance contributing to the positive class. The AM strategy boosts the derived attention weight of each instance by a constant factor depending on the (prior) expected label of that instance. On the other hand, AT imposes an additional cross entropy loss to minimize the discrepancy between the derived attention weights and the prior distributions of the instance labels.

### 2.2 Modeling cell subpopulations

Instead of directly applying the MIL framework, other works have attempted to model the cell subpopulations in each scRNA-seq sample and derive their attributes for prediction. CloudPred (He et al., 2021) models the distribution of cells with a Gaussian mixture model (GMM) and uses the prevalence of the Gaussian clusters as features for prediction. Instead of directly assigning cells to clusters or assuming a distribution of cells, ProtoCell4P (Xiong, Bekiranov, and Zhang, 2023) directly maintains the representations of cell subpopulations, termed prototypes, in the latent space and regularizes the latent embeddings of cells to cluster around the prototypes. In CloudPred, the parameters of the GMM are initialized with the expectation-maximization (EM) method, while the prototypes in ProtoCell4P are initialized randomly. Both frameworks are then optimized end-to-end with respect to the prediction loss via stochastic gradient descent.

## 3 Methodology

### 3.1 Overview

We present a novel approach to incorporate hierarchical information into a MIL framework for patient phenotype prediction with scRNA-seq data. Our approach processes gene expressions or latent representations of individual cells and performs a two-step aggregation procedure, first over cells and then over cell types to produce sample-level representations for prediction. Note that the first step involves either mean-pooling or attention-based aggregation of cells, while the second step performs attention-based aggregation of cell types to produce sample representations. For easy reference, the model with attention only on cell types is termed the Cell Type Attention (CTA) model, and the other one with attention on both cells and cell types is termed the Hierarchical Attention (HA) model. See Figure 1 for an illustration of our proposed method.

**Figure 1:**
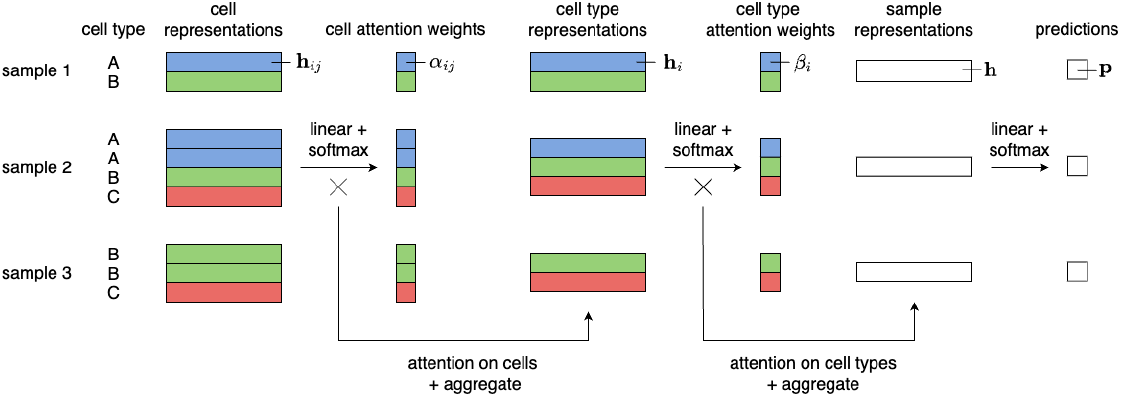
Workflow of the proposed hierarchical attention-based MIL framework for patient phenotype classification using single-cell RNA sequencing (scRNA-seq) data. The model aggregates cell-level representations and applies attention weights at both the cell and cell type levels to generate a sample-level representation for classification. Note that the derivation of the cell attention weights *α*_*ij*_ in the illustrated workflow is specific for the HA model. For the CTA model, the cell attention weights *α*_*ij*_ are equal for all cells in the same cell type (see Eq. 4).

We show that the final prediction can be decomposed into a sum of either cell or cell type contributions, hence providing interpretability at both levels. Based on this, we derive a metric termed importance score that measures the overall contribution of each cell type to the prediction. Moreover, we propose using a permutation test to determine cell types with a significant importance score, which are the ones that might play a role in the mechanism of the patient phenotypes.

### 3.2 Model Architecture

We assume to have data from *S* samples. Data from sample *s* ∈ {1, … , *S*} is denoted as (*D*_*s*_, *y*_*s*_), where *y*_*s*_ ∈ 𝒴 = {1, … , *C*} is the class label, and *D*_*s*_ denotes the single cell data. We assume that single cells from each sample are annotated with cell type label *i* ∈ {1, … , *I* }, where *I* denotes the number of cell types. These annotations can be obtained with any existing cell type annotation tool, or can be manually labeled. For each sample *s*, single cell data *D*_*s*_ consists of gene expression profiles or (possibly lower-dimensional) representations of individual cells and is denoted as *D*_*s*_ = {**x**_*sij*_}, where **x**_*sij*_ ∈ ℝ^*m*^ is the expression profile of the cell *j* = 1, … , *n*_*si*_ with cell type *i* in sample *s* (or 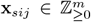 in the case of original gene expression counts). Sample *s* contains altogether 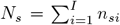 individual cells. Also note that sample *s* does not necessarily contain all *I* cell types, i.e., *n*_*si*_ can equal zero for some *i*. For the clarity of notation, we drop the sample index *s* below and present our method for a single sample, i.e., the single cell data of a single sample is denoted as *D* = {**x**_*ij*_}.

#### 3.2.1 Low-dimensional representations

Before applying the attention mechanism, we pass the input *D* = {**x**_*ij*_} through feed-forward neural network layers (with either one or two hidden layers) to obtain a lower-dimensional representation of each cell, denoted as **h**_*ij*_ ∈ ℝ^*d*^. For example, for a single hidden layer model, the cell representations are computed using the standard feed-forward model

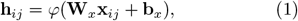

where *φ*(·) is a chosen non-linearity, such as ReLU, and **W**_*x*_ ∈ ℝ^*d*×*m*^ and **b**_*x*_ ∈ ℝ^*m*^ are trainable parameters. These cell representations serve as input for all subsequent layers of the neural network, including the two aggregation modules and the classification layer that are described next.

#### 3.2.2 Cell-Level Aggregation

In this step, the HA model performs an attention-based aggregation over cells. Specifically, for each cell *j* within cell type *i*, we compute a cell-level attention score *α*_*ij*_ using a linear transformation followed by a softmax function within the cell type:

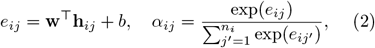

where **w** ∈ ℝ^*d*^ and *b* ∈ ℝ are learnable parameters shared across all cell types, and *α*_*ij*_ is normalized over the cells within cell type *i*. For each cell type *i*, we then compute an auxiliary cell type representation **h**_*i*_ by aggregating the representations of its cells using the cell-level attention weights:

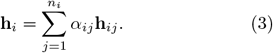

Alternatively, in the CTA model, the cell type representation **h**_*i*_ is obtained via mean-pooling of the cell representations **h**_*ij*_:

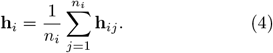

The factor *α*_*ij*_ is therefore defined as 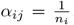. The CTA model effectively simplifies the cell type representation computation by using equal weights instead of the cell-specific attention weights.

Note that this step is the only difference between the CTA and the HA models. All previous and subsequent steps are described jointly for the two models.

#### 3.2.3 Cell Type-Level Aggregation

At the second level of hierarchy, we compute a cell type-level attention score *β*_*i*_ using a linear transformation followed by a softmax function over the cell types:

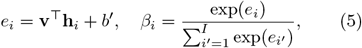

where **v** ∈ ℝ^*d*^ and *b*^*′*^ ∈ ℝ are learnable parameters, and *β*_*i*_ is normalized over the cell types within a sample.

#### 3.2.4 Sample-Level Representation

For each sample, we define a sample-level representation by aggregating the auxiliary cell type-level representations using the cell type-level attention weights:

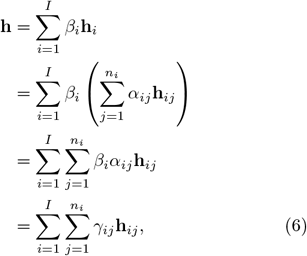

where we have used Eq. (3), and defined

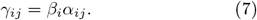

Eqs. (6-7) demonstrates that our definition of sample-level representation results in a hierarchical two-step aggregation mechanism, first over the cell representations to obtain the cell type representations, and then over the cell type representations to obtain the sample-level representations. Note that our hierarchical aggregation results in normalized attention weights since Σ_*i*_ Σ_*j*_ *γ*_*ij*_ = Σ_*i*_ *β*_*i*_ Σ_*j*_ *α*_*ij*_ = _*i*_ *β*_*i*_ = 1.

#### 3.2.5 Classification Layer

A final linear layer maps the representation of each sample to probabilities **p** = [*p*_1_, … , *p*_*C*_]^⊤^ of the *C* classes:

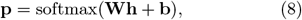

where **W** ∈ ℝ^*C*×*d*^ and **b** ∈ ℝ^*C*^ are learnable parameters shared across all samples.

The hierarchical attention-based MIL model described above is applied separately to each of the *S* samples to obtain sample-specific prediction probabilities **p**_*s*_ = [*p*_*s*1_, … , *p*_*sC*_ ]^⊤^. To define a training objective, we employ the multi-class cross-entropy loss:

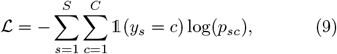

where 𝟙(·) is the indicator function. The objective in Eq. (9) is minimized wrt. model parameters (**W**_*x*_, **b**_*x*_, **w**, *b*, **v**, *b*^*′*^, **W, b**) using gradient descent.

### 3.3 Model Interpretability

Following Engelmann et al., 2024, we show that since the classification layer and the pooling function are both linear, the logits of the final prediction readily decompose into a sum of either cell-level or cell type-level contributions.

Let **Wh** = **z** in Eq. (8), then **p** = softmax(**z** + **b**). Using Eq. (6) we observe that

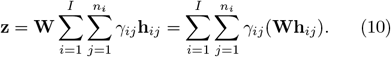

This decomposition highlights that the final logits softmax^−1^(**p**) = **z** + **b** are a sum of the cell-level logits *γ*_*ij*_**Wh**_*ij*_, shifted by the bias **b**. Similarly,

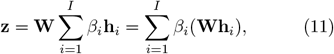

showing that the final logits

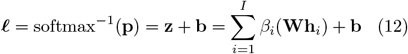

are a sum of the cell type-level logits *β*_*i*_**Wh**_*i*_, shifted by the bias **b**.

### 3.4 Identifying Critical Cell Types

As our models are interpretable at the cell type level, we can identify the *critical cell types* that are most relevant for the prediction and thus likely plays a role in the expression mechanism of the disease phenotypes or clinical outcomes.

To identify these critical cell types, we first decompose the prediction of each sample into a sum of cell type-level logits. In the case of binary classification, a positive cell type logit indicates contribution to the positive class, while a negative logit represents contribution to the negative class. Hence, on average, cell types with higher logits in positive samples and lower logits in negative samples contribute more to the correct prediction. We introduce a metric termed *importance score* to measure the overall contribution of each cell type:

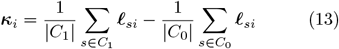

where ***κ***_*i*_ is the importance score of cell type *i, C*_0_ is the set of class 0 samples, *C*_1_ is the set of class 1 samples, and **𝓁**_*si*_ = *β*_*i*_**Wh**_*si*_ is the logit of cell type *i* in sample *s* following the decomposition demonstrated in Eq. 11.

We suggest using a permutation test to identify cell types with a statistically significant importance score. Specifically, we repeatedly permute the sample labels, train the model on this permuted data, and compute the cell type importance scores. This results in a null distribution of importance scores for each cell type, from which we can calculate the empirical *p*-values and identify the critical cell types. Note that since we are essentially performing one statistical test for each cell type, multiple testing correction is needed to avoid inflating the false positive rate.

## 4 Experiments and Results

### 4.1 Data

We evaluate our proposed model on three scRNA-seq datasets: Cardio (Chaffin, Papangeli, Simonson, et al., 2022), COVID (Ziegler, Miao, Owings, et al., 2021), and ICB (Gondal, Cieslik, and Chinnaiyan, 2024). The key characteristics of each dataset are given in Table 1.

**Table 1:**
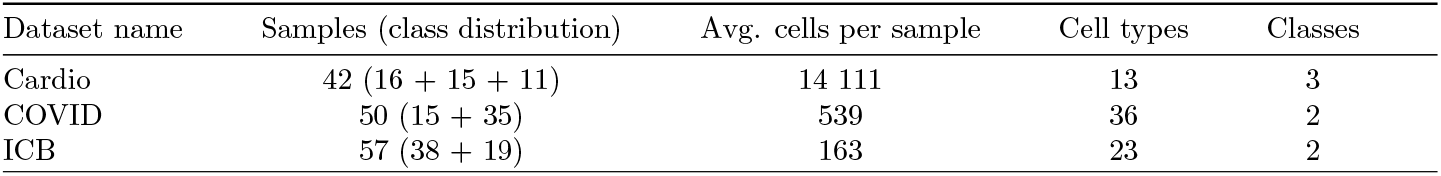
Summary of datasets.

| Dataset name | Samples (class distribution) | Avg. cells per sample | Cell types | Classes |
| --- | --- | --- | --- | --- |
| Cardio | 42 (16 + 15 + 11) | 14 111 | 13 | 3 |
| COVID | 50 (15 + 35) | 539 | 36 | 2 |
| ICB | 57 (38 + 19) | 163 | 23 | 2 |

The Cardio dataset published by Chaffin, Papangeli, Simonson, et al., 2022 contains single-nucleus expression profiles of patients with dilated and hypertrophic cardiomyopathy. The task we perform on this dataset is to classify the samples based on the patients’ disease status, either dilated cardiomyopathy, hypertrophic cardiomyopathy, or normal (healthy control). Following Xiong, Bekiranov, and Zhang, 2023, to preprocess the data, we removed genes with nonzero expressions in fewer than 5 cells, normalized the total gene expression counts of each cell to sum up to 10^4^, and log-transformed the counts. We then used the scGPT model (Cui, Wang, Maan, et al., 2024) with pretrained weights from the whole-human checkpoint to extract the embedding for each cell. Each cell is thus represented by a vector of *m* = 512 dimensions, which serves as input to the model.

The COVID dataset published by Ziegler, Miao, Owings, et al., 2021 provides single-cell expression profiles of COVID-19 patients with three disease statuses (COVID-19, long COVID-19, respiratory failure) and of healthy controls. Since the long COVID-19 and respiratory failure classes contain too few samples, samples from these two classes are removed, transforming the task into binary classification. Following Xiong, Bekiranov, and Zhang, 2023, the preprocessing steps involve removing genes with nonzero expressions in fewer than 5 cells, normalizing the total gene expression counts of each cell to sum up to 10^4^, and log-transforming the counts. We then annotate the single cells using the R package singler (Aran, Looney, Liu, et al., 2019) with the Human Primary Cell Atlas reference dataset (Mabbott et al., 2013). The scGPT model (Cui, Wang, Maan, et al., 2024) with pretrained weights from the whole-human checkpoint is then used to embed cells into a *m* = 512 dimensional vector space.

The ICB dataset is a pan-cancer dataset on patients undergoing the immune checkpoint blockade (ICB) treatment. The dataset consists of scRNA-seq samples from eight studies, compiled and preprocessed by Gondal, Cieslik, and Chinnaiyan, 2024. We are using pre-treatment samples from three studies (Alvarez-Breckenridge, Markson, Stocking, et al., 2022; Pozniak, Pedri, Landeloos, et al., 2024; Bassez, Vos, Van Dyck, et al., 2021) and two cancer types (melanoma and breast cancer) with different subtypes. The objective is to predict whether patients response favourably (class 1) or unfavourably (class 0) to ICB treatment. The samples are preprocessed and downsampled to at most 200 cells per sample by Gondal, Cieslik, and Chinnaiyan, 2024. Since the original data does not contain cell type annotations, we annotated the cells using the R package singler (Aran, Looney, Liu, et al., 2019) with the Blueprint/ENCODE reference dataset (The ENCODE Project Consortium, 2012; Martens and Stunnenberg, 2013). For this task, we use a set of genes which is a merger of predictive/immune gene signatures reported in previous studies. Genes missing from each dataset are removed, resulting in a set of *m* = 824 genes. The normalized expressions of these genes serve as input to the model. The list of gene signatures is given in Section 1 of Supplementary Information.

### 4.2 Baselines and Evaluation

We benchmark our proposed models against four existing methods: ScRAT (Mao, Lin, Wong, et al., 2024), ProtoCell4P (Xiong, Bekiranov, and Zhang, 2023), CloudPred (He et al., 2021), and MixMIL (Engelmann et al., 2024). All models are evaluated following a repeated, nested 10-fold cross validation (CV) procedure. Specifically, we perform nested CV and calculate the AUC on the stacked predictions from all folds. The hyperparameters of our models and competing models are optimized using the inner CV loop, such that the combination of hyperparameters that achieves the highest AUC on the stacked predictions from all inner CV folds is used to train the model. This procedure is repeated 10 times with different random seeds for CV splitting, resulting in 10 AUCs, and the mean and standard deviation of the AUCs are reported. For more details on hyperparameter tuning, please see Section 2 of Supplementary Information.

The evaluation results of all six models on the three benchmark datasets are given in Table 2. Both of our models consistently outperform existing models on all datasets, demonstrating the utility of our proposed framework. It is worth noting that on the COVID dataset, when using the original annotations by Ziegler, Miao, Owings, et al., 2021 instead of the ones obtained with singler, ProtoCell4P achieves an AUC of 0.89 ± 0.02, which is on par with our proposed models.

**Table 2:** Comparison of AUC scores across different models and datasets. The two best mean AUCs for each dataset are highlighted in bold.

| Model | Cardio | COVID | ICB |
| --- | --- | --- | --- |
| ScRAT | $0.85 \pm 0.02$ | $0.83 \pm 0.04$ | $0.70 \pm 0.06$ |
| ProtoCell4P | $0.91 \pm 0.02$ | $0.85 \pm 0.02$ | $0.68 \pm 0.04$ |
| CloudPred | $0.82 \pm 0.02$ | $0.82 \pm 0.05$ | $0.58 \pm 0.05$ |
| MixMIL | $0.87 \pm 0.02$ | $0.83 \pm 0.03$ | $0.65 \pm 0.06$ |
| Our model (CTA) | **$0.98 \pm 0.01$** | **$0.90 \pm 0.03$** | **$0.77 \pm 0.04$** |
| Our model (HA) | **$0.99 \pm 0.02$** | **$0.89 \pm 0.03$** | **$0.75 \pm 0.06$** |

Also note that our two models, CTA and HA, achieve comparable performance on all benchmark datasets. As the CTA model involves a mean-pooling over cells, it assumes equal contribution of cells towards the cell type representation, which proves to be adequate for the tasks at hand. However, on more complex datasets with heterogeneous cell subpopulations in the same cell type, we postulate that the HA model might be more suitable, as it accounts for the variability of both cells and cell types in predicting the label.

### 4.3 Impact of Data Quality

Sample sizes are typically small in early-stage clinical trials, and biological sampling can result in a limited number of single cells. Moreover, our method relies on existing cell type annotations or cell type annotation methods, which can be inaccurate. To evaluate the robustness of our model against these variations in data quality, we conduct three additional experiments.

#### 4.3.1 Varying Train Size

First, we investigate the impact of limited training size on model performance by sequentially using 25%, 50%, and 75% of the samples for training and reserving the rest for testing. For each training size, we run the experiment for 100 random train-test splits. The AUC is calculated on the testing set at each run, and the mean AUC over 100 runs is reported. This experiment is conducted for all models on all datasets. The results on the Cardio dataset are summarized in Figure 2a. Results on other datasets are given in Figure 1 of Supplementary Information.

**Figure 2:**
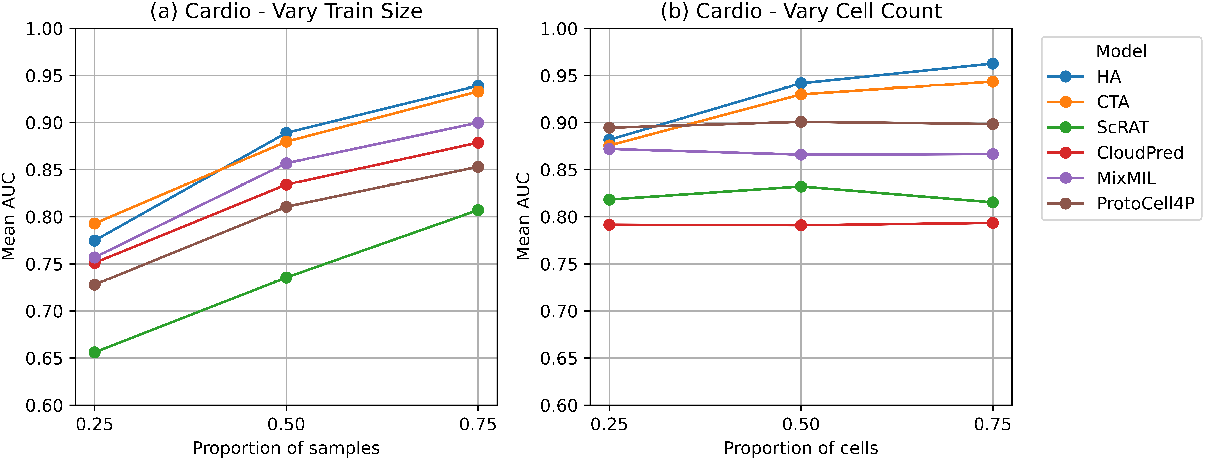
**(a)** Performance of models on the Cardio dataset when the training sizes are varied. *x*-axis corresponds to the proportion of samples in the original dataset, which for the Cardio dataset is *S* = 42; **(b)** Performance of models on the Cardio dataset when the cell counts are varied. *x*-axis corresponds to the proportion of cells in each sample.

Overall, both of our models demonstrate competitive performance, outperforming existing models across all training sizes on the Cardio and ICB datasets. Notably, our models are capable of achieving reasonable AUCs even when trained on only a handful of samples, showing robustness against limited sample sizes.

#### 4.3.2 Varying Cell Counts

The second experiment studies the effect of varying cell counts on performance. We randomly subsample 25%, 50%, and 75% of the cells in each sample and evaluate the model following the repeated, nested CV procedure described in Section 4.2. This process is repeated 10 times for random cell subsampling and CV splitting, resulting in 10 AUCs, and the mean AUC is reported. This experiment is conducted for all models on all except the ICB dataset, since this dataset is already downsampled to at most 200 cells per sample. The results on the Cardio dataset are illustrated in Figure 2b. The results on the COVID dataset are given in Figure 2 of Supplementary Information.

In general, the performance of our models improves as the proportion of cells in each sample increases, suggesting the ability to effectively utilize the information from additional cells. Moreover, comparing the results of the first and second experiment, all tested models appear less sensitive to the number of cells than to the number of training samples. In fact, the performance of existing methods on both benchmark datasets either plateaus or improves slightly as the proportion of cells increases. These results suggest that the performance of existing methods is generally stable after a certain number of cells are included, consistent with the findings of He et al., 2021 .

#### 4.3.3 Randomizing Cell Type Annotations

In the third experiment, we assess the influence of noisy cell type annotations on the performance of our models. We sample 25% and 50% of cells in each sample and assign them random cell types drawn from a uniform distribution over all cell types in the dataset. For each ratio of noisy annotations, we evaluate our models using the nested CV procedure described in Section 4.2. This process is repeated 10 times for random cell sampling and annotation reassignment, resulting in 10 AUCs, and the mean AUC is reported. Note that this experiment is only conducted for our models, since other models do not require cell type annotations as input (except for ProtoCell4P, which optionally uses cell type annotations). The results on the Cardio dataset are summarized in Figure 3. Results on other datasets are given in Figure 3 of Supplementary Information. Note that the mean AUCs at 0.0 (no random annotations) are the same as the cross validation results presented in Table 2.

**Figure 3:**
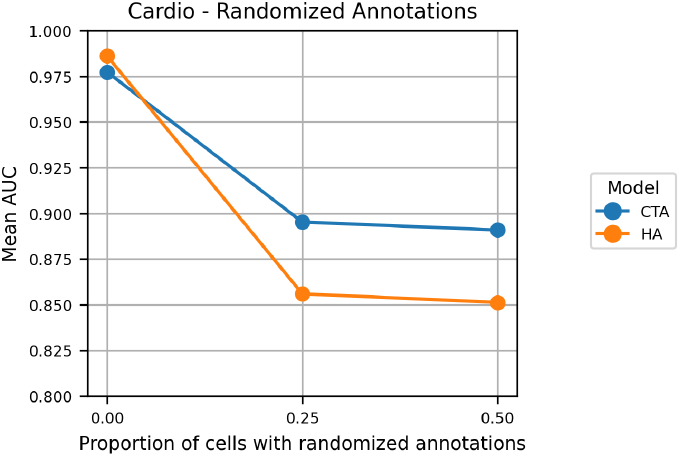
Performance of the proposed models on the Cardio dataset with varying levels of cell type annotation accuracy. *x*-axis corresponds to the proportion of cells that are artificially assigned a randomly chosen cell type annotation.

As our approach relies on existing cell type annotations, the performance of our model declines as the cell type annotations become less accurate. The CTA model appears less sensitive to the quality of the annotations than the HA model, as the mean-pooling operation at the cell level might be more robust to noise compared to the attention-based aggregation operation (Engelmann et al., 2024). This dependence on accurate annotations is a potential downside of our approach and can possibly be alleviated with data augmentation strategies. However, results obtained with off-the-shelf cell type annotation tools, such as singler (Aran, Looney, Liu, et al., 2019), reported in Table 2 still demonstrate highly competitive performance.

### 4.4 Ablation Test

To validate the benefits of incorporating cell type information into the MIL framework, we conduct an ablation test. Specifically, we compare the performance of our two proposed models with two modified models, one with mean pooling over the cells, and the other with attention on cells. Note that the two modified models do not use cell type annotations, thus allowing us to pinpoint the impact of additionally using this information, either through attention on cell types (CTA model) or through attention on both cells and cell types (HA model).

In the mean-pooling model, the entire attention-based aggregation module is omitted, and the sample-level representation is generated by averaging the celllevel representations:

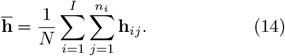

Here, **h**_*ij*_ is the representation of cell *j* of cell type *i, N* is the total number of cells in the sample, and 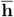 is the sample representation. This corresponds to converting scRNA-seq data to pseudo-bulk representation.

On the other hand, in the cell-attention model, only the cell-type attention layer is omitted, and the sample-level representation is a weighted sum of the cell-level representations:

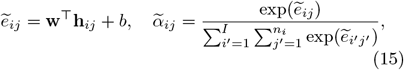

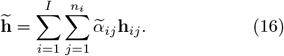

Here, *e*_*ij*_ and *α*_*ij*_ are the unnormalized and normalized attention weights of cell *j* of cell type *i*. This ablation corresponds to the attention mechanism used in MixMIL method (Engelmann et al., 2024).

The two modified models (mean-pooling and cell-attention) are assessed on all three datasets following the repeated, nested cross validation procedure described in Section 4.2.

The results of the ablation tests are summarized in Table 3. Our proposed models outperform the two modified models on all except the COVID dataset, where the mean-pooling model also achieves the highest AUC of 0.90 ± 0.02, the same as the CTA model. This result underscores the utility of incorporating cell type information into the attention-based MIL framework for modeling scRNA-seq data. Notably, the performance edge of the cell type attention (CTA) model over the cell attention model suggests the benefits of applying the attention mechanism on cell types instead of cells.

**Table 3:** Ablation test results. Note that the results for our two proposed models are the same as in Table 2, also presented below for easy reference.. The two best mean AUCs for each dataset are highlighted in bold.

| Model | Cardio | COVID | ICB |
| --- | --- | --- | --- |
| **Mean-pooling** | 0.91 ± 0.02 | **0.90 ± 0.02** | 0.74 ± 0.03 |
| **Cell-attention** | 0.91 ± 0.03 | 0.82 ± 0.04 | 0.70 ± 0.04 |
| **Our model (CTA)** | **0.98 ± 0.01** | **0.90 ± 0.03** | **0.77 ± 0.04** |
| **Our model (HA)** | **0.99 ± 0.02** | 0.89 ± 0.03 | **0.75 ± 0.06** |

One surprising result is that the mean-pooling model performs relatively better than the cell-attention model on all datasets. One possible explanation for this could be that the mean pooling operation offers reduced complexity, which helps with the small sample size and the low signal-to-noise ratio of scRNA-seq data. Also note that the cell-attention model does not equate to the MixMIL model. Despite having the same attention mechanism applied on cells, MixMIL lacks the dimensionality reduction layers and uses a variational inference framework.

### 4.5 Biological Relevance

To investigate the interpretability of our approach, we identify the cell types that are critical for the HA model in the COVID dataset following the permutation testing procedure described in Section 3.4. Moreover, we validate these critical cell types against existing literature, thereby assessing the model’s ability to capture biologically relevant associations. Note that in this experiment, we use the original cell type annotations by Ziegler, Miao, Owings, et al., 2021 as they are more fine-grained than the annotations obtained with singler. For more details on the implementation of the permutation test, please see Section 3 of Supplementary Information.

Table 4 provides the cell type importance scores and their adjusted *p*-values. At the significance level of 0.05, our model identifies 11 cell types as critical in distinguishing between normal and COVID samples: macrophages, ciliated cells, secretory cels, B cells, goblet cells, developing secretory and goblet cells, plasmacytoid DCs (pDCs), basal cells, T cells, squamous cells, and deuterosomal cells. These findings generally align with the disease mechanism of COVID-19 and the biological changes in response to infection.

**Table 4:** Cell type importance scores and adjusted *p*values in the COVID dataset. The critical cell types with significant adjusted *p*-values (*p* ≤ 0.05) are highlighted in bold.

| Cell Type | Importance Score | Adjusted $p$-value |
| --- | --- | --- |
| **Macrophages** | 0.17 | **0.05** |
| **Ciliated Cells** | 0.13 | **0.00** |
| **Secretory Cells** | 0.11 | **0.00** |
| **B Cells** | 0.09 | **0.05** |
| **Goblet Cells** | 0.06 | **0.00** |
| Developing Ciliated Cells | 0.06 | 0.07 |
| **Developing Secretory and Goblet Cells** | 0.06 | **0.05** |
| **Plasmacytoid DCs** | 0.05 | **0.05** |
| **Basal Cells** | 0.05 | **0.05** |
| Mitotic Basal Cells | 0.05 | 0.06 |
| **T Cells** | 0.05 | **0.05** |
| Ionocytes | 0.05 | 0.06 |
| **Squamous Cells** | 0.04 | **0.05** |
| Dendritic Cells | 0.04 | 0.09 |
| **Deuterosomal Cells** | 0.04 | **0.05** |
| Erythroblasts | 0.02 | 0.06 |
| Mast Cells | 0.01 | 0.14 |
| Enteroendocrine Cells | -0.02 | 0.64 |

At the onset of COVID-19, ciliated cells in the nasal respiratory epithelium are known to be the primary sites of viral replication (Ahn, Kim, Hong, et al., 2021). Infection in these cells can damage the ciliated layer and impair the mucociliary clearance function (Robinot, Hubert, De Melo, et al., 2021). Upon the loss of the ciliated layer, basal cells undergo proliferation and differentiation into ciliated cells as part of the repair mechanism of the epithelium. There has been some evidence indicating that secretory and goblet cells are also infection sites, though less prominent than ciliated cells (Zhu, Wang, Liu, et al., 2020; Robinot, Hubert, De Melo, et al., 2021). The significance of the immune cells (macrophages, B cells, T cells, pDCs) might reflect their over-activation and possible exhaustion during the immune response to infection (Alahdal and Elkord, 2022). Among these, pDCs are considered essential during antiviral immune defense (Ngo et al., 2024), serving as the primary antigen presenting cells and regulating responses to SARSCoV infection (Alahdal and Elkord, 2022).

## 5 Discussion and Conclusion

In this work, we introduce two strategies to incorporate hierarchical information into the attention-based MIL framework for patient phenotype classification with scRNA-seq data. Specifically, we propose using an attention mechanism that either operates on cell types or both cells and cell types, with the latter accounting for the heterogeneity of both cells and cell types in contributing to the prediction. Through extensive experiments, we demonstrate the robustness and utility of our proposed models, even in data-constrained scenarios. Interestingly, empirical results show that applying the same attention mechanism on cell types instead of cells improves predictive performance, which validates the benefits of utilizing a hierarchical structure. Our approach provides interpretability at both the cell and cell type levels, thus supporting biological discoveries.

As a possible area for improvement, we note that our attention mechanism is relatively simple. Even though this design choice suits the typically small sample size of real-world scRNA-seq datasets, for larger datasets, more complex attention mechanisms (e.g. the attention mechanism in transformers) could be used to enhance the modeling capacity. We also note that our approach can be easily adapted to account for patient-level covariates, simply by extending the final classification layer to accommodate the additional inputs.

## 6 Code Availability

Our implementation is available at https://github.com/minhchaudo/hier-mil.

## 7 Acknowledgements

We would like to acknowledge the computational resources provided by the Aalto Science-IT.

