## Supplementary Information for "Incorporating Hierarchical Information into Multiple Instance Learning for Patient Phenotype Prediction with scRNA-seq Data"

December, 2024

#### 1 Genes for ICB Treatment Response Prediction

The list of genes used in the ICB dataset is a merger of multiple gene signatures reported in previous studies. These include:

- Stem.Sig (Zhang et al., 2022)
- IFN-gamma and expanded immune signature (Ayers et al., 2017)
- LRRC15-CAF signature (Dominguez et al., 2020)
- Cytolytic activity signature (Rooney et al., 2015)
- NLPR3 inflammasome signature (Ju et al., 2021)
- TIR signature (Mei et al., 2021)
- TIME signature (Lee et al., 2020)
- Immune signatures (HLA (Liu et al., 2018), TILs (Massink et al., 2015), and IFN response(Yoshihara et al., 2013))
- Immune checkpoint genes (Ju et al., 2021)

#### 2 Details on Hyperparameter Tuning

Hyperparameter tuning is performed using cross validation. Specifically, for each pair of training and testing datasets, we perform cross validation on the training data with different combinations of hyperparameters. The AUC is calculated on the stacked predictions from the cross validation folds, resulting in one AUC value for each hyperparameter combination. We use the hyperparameter combination with the highest AUC to re-train the model on the entire training set, and this final model is used for prediction on the test set.

The hyperparameter search space for each model is given below:

- ScRAT:
  - Learning rate: {1e-4, 1e-3, 1e-2}
  - Number of epochs: {100}
  - Number of attention heads: {1, 2, 4}
  - Dropout rate: {0.0, 0.3, 0.5, 0.7}
  - Weight decay: {1e-4, 1e-3, 1e-2}
  - Whether data augmentaion is performed: {True, False}
  - Embedding dimension: {8, 32, 64}
  - Number of augmented samples: {100}
  - PCA: {False}
- ProtoCell4P:

- Learning rate: {1e-4, 1e-3, 1e-2}
- Number of epochs: {50, 75, 100}
- Output dimension of linear layers in encoder and decoder: {32, 64, 128}
- Hidden dimension: {8, 16, 32}
- Number of prototypes: {8, 16, 32}
- Number of epochs in pretraining: {75}

- CloudPred:
  - Learning rate in pretraining: {1e-2}
  - Number of epochs in pretraining: {100}
  - Learning rate: {1e-2, 1e-3, 1e-4}
  - Number of epochs: {100}
  - Number of centers: {2, 8, 16}
- MixMIL:
  - Number of epochs: {100, 500, 1000}
  - Learning rate: {1e-2, 1e-3, 1e-4}
- Our models (CTA and HA):
  - Number of epochs: {100, 500, 1000}
  - Dropout rate: {0.0, 0.3, 0.5, 0.7}
  - Weight decay: {1e-4, 1e-3, 1e-2}
  - Hidden dimensions: {32, 64, 128}
  - Number of linear input layers: {1, 2}
  - Learning rate: {5e-3, 1e-3}

Other hyperparameters are kept at their default values.

Since the hyperparameter search spaces of our models, ScRAT, and ProtoCell4P are large, we perform Bayesian hyperparameter tuning with the Python package `optuna` (Akiba et al., 2019). The number of trials (number of hyperparameter combinations to try) is set to 30. For other models, since the search space is relatively small, grid search is used.

### 3 Implementation of the Permutation Test

To compute the cell type importance scores, we first perform 5-fold cross validation and stack the predictions from all folds to obtain the predictions for the entire dataset. The importance score for each cell type can then be calculated from the predictions by using Eq. 13 in the main text. The null distribution of importance scores is obtained with 100 permutations. Multiple testing correction is performed with the Benjamini-Hochberg method.

### 4 Results of Experiments on Varying Data Quality

Figure 4 illustrates the performance of all models when the proportion of training samples is varied. Figure 4 shows the performance of all models with various configurations of cell counts. Figure 4 summarizes the performance of our proposed models when cell type annotations are randomized.

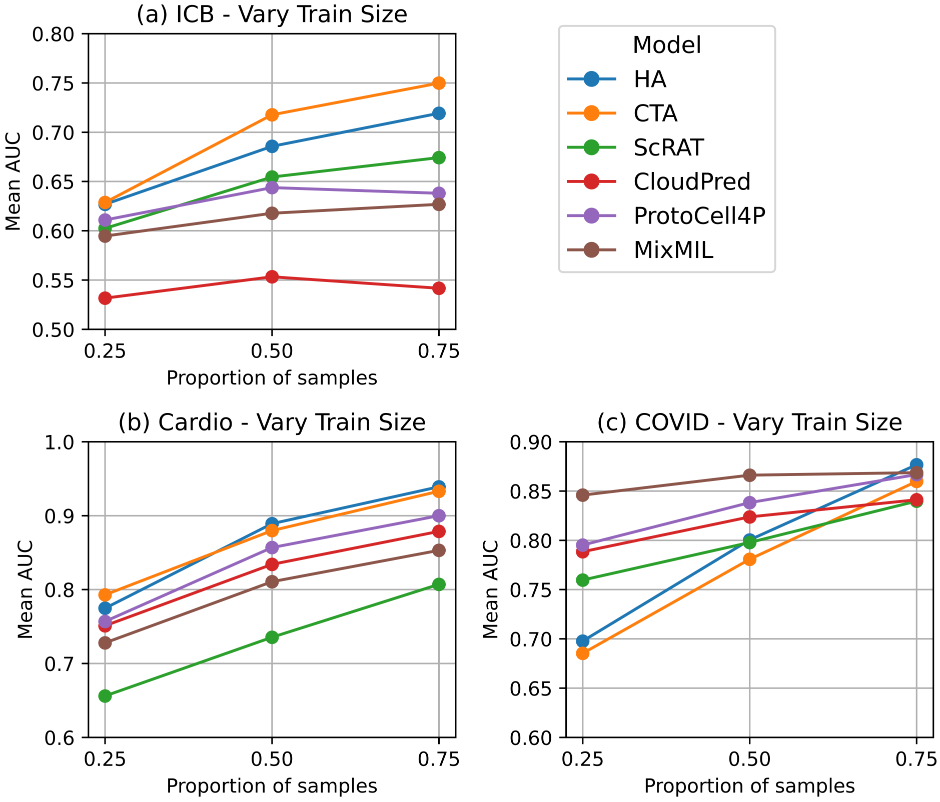


Figure 1: Performance of proposed and existing models on three datasets ((a) - ICB, (b) - Cardio, (c) - COVID) across different proportions of training samples.

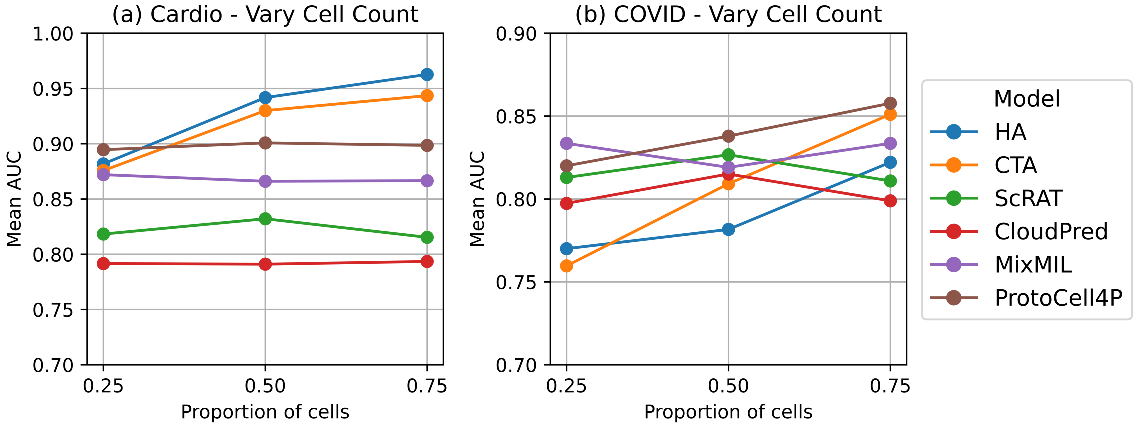


Figure 2: Performance of proposed and existing models on two datasets ((a) - Cardio, (b) - COVID)) across different proportions of cells in each sample.

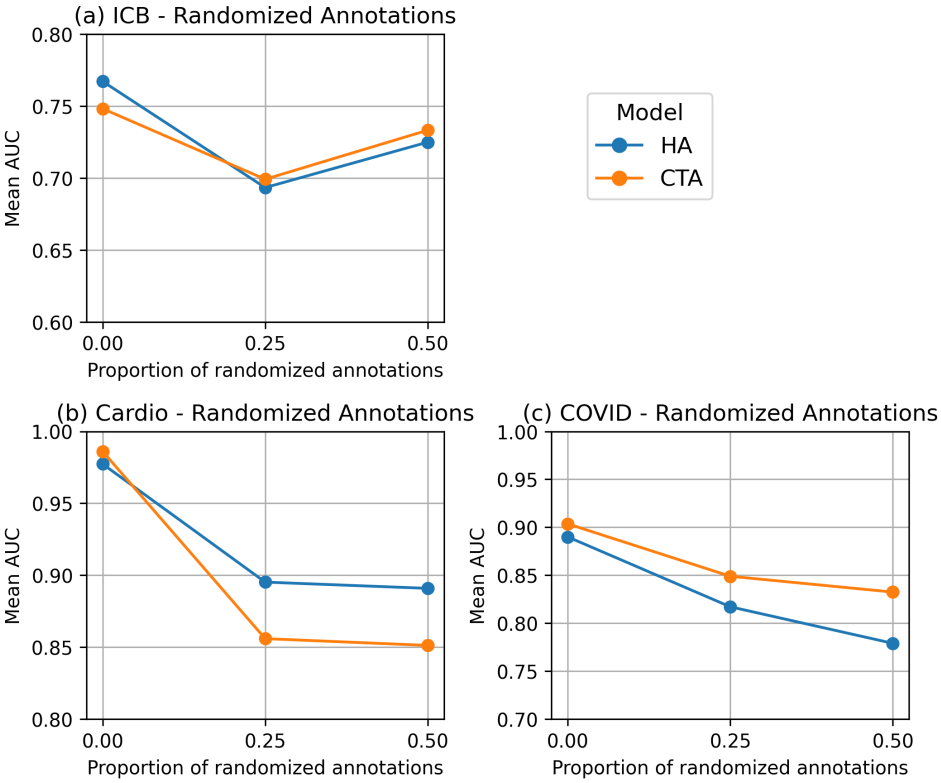


Figure 3: Performance of proposed models on three datasets ((a) - ICB, (b) - Cardio, (c) - COVID) across different proportions of randomized cell type annotations.
